# Sympathetic Controls Fate and Function of Adult Sensory Neurons

**DOI:** 10.64898/2026.02.12.702354

**Authors:** Debora Denardin Lückemeyer, Wenrui Xie, Arthur Silveira Prudente, Judith A. Strong, Temugin Berta, Jun-Ming Zhang

## Abstract

Sensory neurons in the dorsal root ganglia (DRG) serve as conduits for transmitting peripheral stimuli to the central nervous system, playing an essential role in sensory perception and coordinated movement. This study reveals that sympathetic innervation is critical for these neurons’ survival and functional integrity. Using a surgical microsympathectomy model in mice, we found that targeted sympathetic denervation of the lumbar DRGs triggered robust neuronal death, peaking four days post-surgery. This cellular loss is evidenced by reduced spinal projections, decreased nerve density in the skin, and impaired sensory and motor functions. We further identified norepinephrine (NE) as a vital neuroprotective agent; continuous NE supplementation effectively prevented cell death. Additionally, macrophage activation following denervation proved protective, as macrophage depletion exacerbated neuronal loss. It is suggested that the loss of sympathetic input disrupted mitochondrial homeostasis, releasing Smac/DIABLO, activating Caspase-3, and leading to cell death. These findings highlight the sympathetic nervous system’s novel role in maintaining sensory neuron viability through tonic adrenergic support.

## Introduction

Sensory neurons such as dorsal root ganglion (DRG) neurons and trigeminal ganglion neurons are key components of the somatosensory nervous system. These primary afferent neurons serve as essential conduits for transmitting both external and internal stimuli from periphery to the central nervous system (CNS), thereby playing a significant role in how organisms perceive and react to their environment. The diversity of sensory neuron subtypes is crucial for their ability to distinguish between various sensory modalities, including tactile sensations, thermal stimuli, nociceptive signals for pain, and proprioceptive feedback, which is critical for coordinated movement (*1–3*).

The survival and functionality of adult sensory neurons are influenced by several factors, including neurotrophic support from peripheral sources, cellular signaling processes, and interactions with neighboring non-neuronal cells like resident macrophages and satellite glia. Neurotrophic factors such as nerve growth factor (NGF) and brain-derived neurotrophic factor (BDNF) are instrumental in supporting the health and maintenance of sensory neurons (*4*). These factors, along with glial cell interactions, provide crucial signals that help maintain neuronal integrity (*5*).

The sympathetic nervous system is known for its role in managing physiological responses to stress and maintaining immune homeostasis through mechanisms that include modulation of heart rate and blood flow during physical or perceived stressors (*6–9*). In normal DRGs, tyrosine hydroxylase (TH)-positive sympathetic fibers are mostly observed on the surface of the whole DRG and along arterioles within the ganglion. Under pathological conditions such as nerve injury or inflammation, sympathetic fibers often extend away from the surface and blood vessels in the DRGs, sometimes forming basket-like structures around a subset of sensory neurons (*10–12*). The sympathetic system plays a significant role in modulating sensory neuron activity, regeneration of the injured nerve, and inflammatory responses, all of which are crucial to the development and persistence of chronic pain conditions (*13–20*).

Clinically, interventions such as surgical or chemical sympathectomy, and regional sympathetic block, are utilized to manage pain by targeting conditions like complex regional pain syndrome (CRPS) and other neuropathic and inflammatory conditions (*21–24*). In rodent models, these interventions often result in partial or complete alleviation of pain symptoms by modulating local immune and inflammatory activities (*14–18, 25*).The primary sympathetic neurotransmitter, norepinephrine (NE), exerts context-dependent effects: it promotes neuronal survival via PI-3K signaling in the CNS (*26, 27*), yet can induce apoptosis in cardiac myocytes through β-adrenergic pathways(*28–30*). Similarly, macrophages in the DRG exhibit dual functions that can be both protective and damaging. While they help clear debris, release neurotrophic factors, and modulate immune responses to support neuronal survival and tissue repair (*31, 32*), activated macrophages can also produce pro-inflammatory cytokines and reactive species that may contribute to neuronal damage, exacerbate neuroinflammation, and promote degenerative processes (*33*). This duality underscores the complex role of sympathetic innervation and macrophages in maintaining DRG health and in the pathology of nerve injury.

Here we report that sympathetic innervation is vital for the survival and functional integrity of primary sensory neurons in otherwise healthy mice. Disruptions in sympathetic innervation of lumbar DRGs result in a significant loss of sensory neurons, which is prevented by supplementing NE. Such neuronal loss following sympathetic denervation is associated with significant functional impairments, affecting both pain perception and sensory information processing. These findings unveil a previously unrecognized physiological role of the sympathetic nervous system and reveal a novel therapeutic mechanism underlying the anti-hyperalgesic effects of sympathetic blockade.

## Results

### Robust cell death in the DRGs following targeted sympathetic denervation

Surgical microsympathectomy (mSYMPX) for targeted sympathetic denervation (TSD) is a well-established technique developed and routinely utilized in our lab (*14, 15, 17, 20, 25*) to explore the functional roles of the sympathetic nervous system in regulating pathological pain and inflammation. By integrating this technique with *in vivo* microscopic calcium imaging, we have been able to directly visualize and record neuronal activity in animals, providing valuable insights into neuronal dynamics both with and without mSYMPX intervention. During our imaging sessions, we observed substantial differences in the size of the DRGs and the number of neurons present in the DRGs of animals underwent mSYMPX. These changes indicate that sympathetic denervation may lead to significant neuronal loss within the DRGs.

We thus investigated the role of the sympathetic signaling in maintaining the cellular health of sensory neurons. To do this, we performed a mSYMPX to selectively denervate L3 and L4 DRGs in otherwise healthy mice. mSYMPX is a minimally invasive procedure that involves transecting the gray ramus communicans innervating individual DRG (Fig. 1A). We injected propidium iodide (PI) intravenously, a fluorescent dye that stains dead cells by entering through compromised cell membranes and binding to DNA (*34–38*) followed by isolation and removal of the sympathetically denervated DRGs for *ex vivo* imaging of sensory neurons (Fig. 1B). Compared to the contralateral side of operated animals, the number of PI+ neurons directly visualized under the microscope was significantly higher in the DRGs with sympathetic denervation on postoperative day (POD) 2 (Fig.1C). The number of PI+ neurons reached the peak on POD4 and returned to the control level by POD7 (Fig. 1C), which suggests that cell death/cell loss only occurred in the very early stage following sympathetic denervation. Although cell death also occurred in some non-neuronal cells, neuronal cells could be easily identified by its visual appearance under microscope (Fig. 1D).

**Figure 1.**
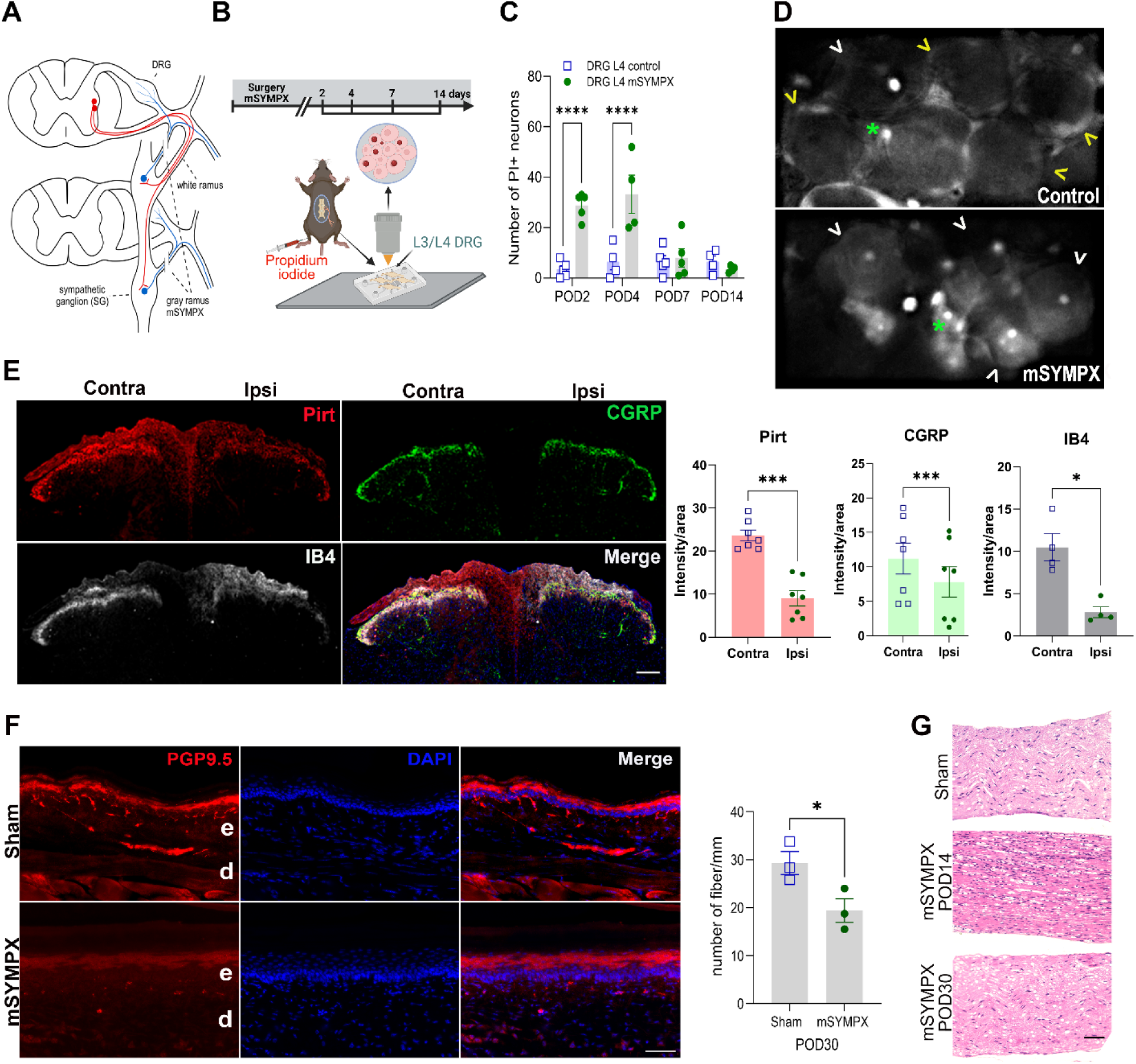
Evidence of sensory neuron death following targeted sympathetic denervation of the lumbar DRGs. **(A)** Schematic figure of the mSYMPX surgery which consisted of cutting the gray rami from the L3 and L4 postganglionic sympathetic ganglia. Blue line indicates a postganglionic sympathetic fiber (gray ramus); Red line indicates the white ramus. **(B)** Diagram showing the timeline and the in vivo recording setup used to count the number of propidium iodide (PI)-positive neurons in the DRG. **(C)** Quantification of PI-positive neurons in dorsal root ganglia (DRG) at 2, 4, 7 and 14 days following mSYMPX. Two-way ANOVA followed by Bonferroni’s multiple comparisons test revealed significant differences between L4 control and mSYMPX groups at POD 2 and POD4 (****p < 0.0001), while differences at POD7 and POD14 were not significant (p > 0.9999), n=4-5 per group. **(D)** Representative images of PI signals in DRG neurons from sham and mSYMPX groups. White arrowheads indicate PI-positive neurons, yellow arrowheads indicate PI-negative neurons, and green asterisks denote other PI-positive cells. **(E)** Representative image of a mouse spinal cord section 30 days post-mSYMPX surgery, immunolabeled for Pirt (phosphoinositide-interacting regulator of TRP channels), CGRP (calcitonin gene-related peptide), and IB4 (isolectin B4), with nuclear counterstaining using DAPI (blue). Scale bar: 100 µm. Images are representative of data obtained from 4–7 mice. On the ipsilateral side of the spinal cord, expression levels of Pirt, CGRP, and IB4 were reduced compared to the contralateral side. Fluorescence intensity values were normalized to the corresponding area. Statistical analysis was performed using two-tailed, paired t-test, revealing significant differences between groups (*p < 0.05, ***p < 0.0001). **(F)** Representative images of paw skin at 30 days after mSYMPX surgery. PGP9.5 immunofluorescence in skin sections, measuring IENF. Sham controls exhibit normal fiber density, while the mSYMPX group shows a significant decrease in fiber numbers after 30 days (N = 3). Quantification of IENF, presented as the number of fibers per linear millimeter of epidermis. Epidermis (e) and dermis (d) are indicated. Scale bar = 50 µm. **(G)** Hematoxylin and eosin (H&E) staining of the contralateral and ipsilateral sciatic nerves, showing increase of immune cell infiltration on POD14 and decreased nerve fiber density in the ipsilateral side after 30 days of mSYMPX. Scale bar = 50 µm.

Next, we examined the spinal projections of sensory neurons in sympathectomized animals and compared the ipsilateral and contralateral side of the spinal cord. Sympathetic denervation resulted in significant cell death and loss of sensory neurons, leading to a marked reduction in spinal projections. The Pirt gene is expressed in all sensory neurons, allowing comprehensive labeling of the sensory neuron population (*39*). As shown in figure 1E, there was a remarkable decrease in spinal projections on the ipsilateral side of the dorsal horn in Pirt-cre mice crossed with tdTomato on POD30. This reduction was further confirmed by the decrease of calcitonin gene-related peptide (CGRP+) and isolectin IB4 (IB4+) positive signals in the dorsal horn, indicating loss of spinal projections from both peptidergic and non-peptidergic sensory neurons. The loss occurred primarily in laminae I–III, with the IB4+ signal in lamina II being particularly affected.

Sensory neuron loss also significantly decreased the density of sensory fibers in the skin, mirroring the reduction in spinal projections. Immunohistochemical analysis using the pan-neuronal marker PGP9.5 revealed reduced nerve fiber density in both the epidermis and dermis of mSYMPX animals at POD30 (Fig. 1F).

We also examined the sciatic nerve in mice with and without mSYMPX on POD14, and 30. H&E staining of the sciatic nerve on POD14 showed increased inflammatory cell infiltration (Fig. 1G), which indicates a potential ongoing degeneration of the nerve fibers from the lost neurons. These results are consistent with the observed reductions in central projection and peripheral innervation of the sensory neurons.

To characterize the lost sensory neurons, we performed IHC of the sympathetically denervated DRGs with antibodies against neurofilament 200 (NF200), IB4 (non-peptidergic), CGRP (peptidergic), or tyrosine hydroxylase (TH - VGLUT3), and compared to the DRGs with intact sympathetic innervation on POD 30 (Fig.2A). Comparing DRGs with intact sympathetic innervation, DRGs with sympathetic denervation showed decreased numbers of CGRP-, TH-, and IB4-positive neurons. The NF200 positive neurons, however, did not show significant alteration between the DRGs with and without microsympathectomy (Fig. 2B).

**Figure 2:**
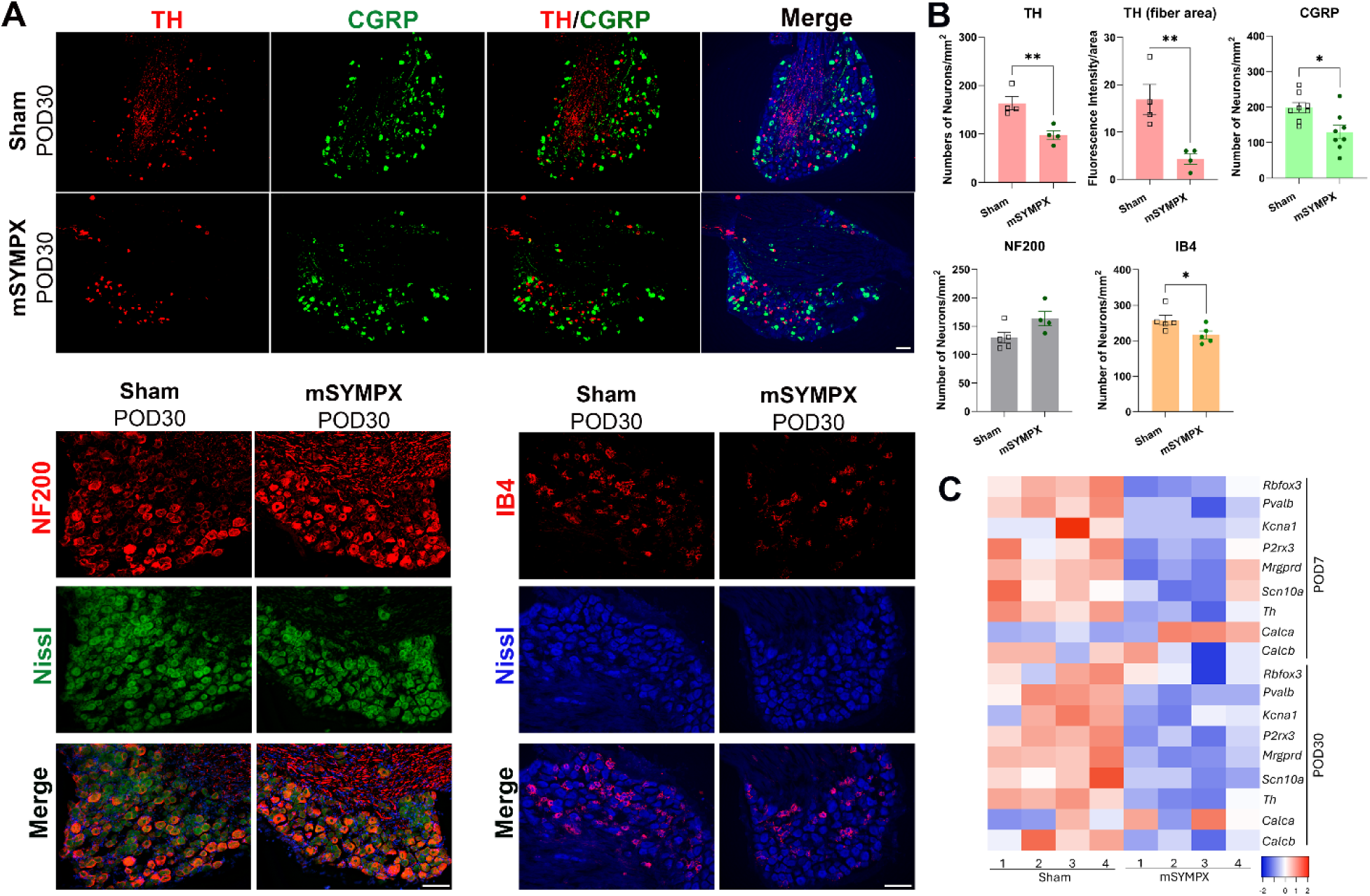
Changes in cellular and gene expressions in the DRGs following sympathetic denervation. **(A)** Immunohistochemistry of tyrosine hydroxylase (TH), calcitonin gene-related peptide (CGRP), neurofilament 200 (NF200), isolectin (IB4), and Nissl staining in sham and mSYMPX mice at POD30. Scale bar: 100 µm. **(B)** Quantification of positively labeled neurons in the cellular region of DRG, showing significant differences between control and mSYMPX animals at POD 30 (n=4-8). Statistical analysis was performed using two-tailed student t-tests, revealing significant differences between groups (*p < 0.05, **p < 0.01). **(C)** qPCR analysis of neuronal gene marker expressions at POD 7 and 30 revealing persistent transcriptional alterations following mSYMPX. The heatmap represents relative gene expressions, with blue indicating downregulation and red indicating upregulation.

Using qPCR, we investigated the expression levels of neuronal markers, including *Rbfox3, Pvalb, Kcna1, P2rx3, Mrgprd, Scn10a, Th, Calca, and Calcb*, in the DRG following sympathetic denervation at POD7 and POD30. Our analysis revealed that all genes examined, except *Calca*, exhibited downregulation, as shown in the heatmap (Fig. 2C and Suppl. Fig. 1). Given that cell death initiated by mSYMX occurred only within the first few days, the changes in gene expression observed at POD7 and POD30 reflect the gene regulation occurring in the surviving DRG neurons. These findings suggest that the effects of sympathetic denervation on sensory neurons were more extensive than anticipated, with some neurons experienced alterations in gene expression while remaining viable.

### Altered sensory functions following sympathetic denervation of the DRGs

Primary sensory neurons detect and transmit specific sensory modalities (e.g., touch, temperature, and pain) by converting physical and chemical stimuli from the peripheral tissues into electrical signals for the central nervous system. We then assessed different sensory modalities following sympathetic denervation of the sensory ganglion.

Initial testing with von Frey filaments did not reveal any changes in mechanical sensitivity on the plantar surface of the paw at POD2 or subsequent time points up to POD28 (Fig.3A). This may reflect the limitations of the von Frey test in detecting changes at forces beyond the highest levels applied. In contrast, pinprick test using sharp needle attached to 1-gram (bending force) nylon filament demonstrated a marked decrease in evoked responses in the sympathectomized mice compared to the sham-operated controls but only on POD7 and POD14 (Fig 3B).

**Figure 3.**
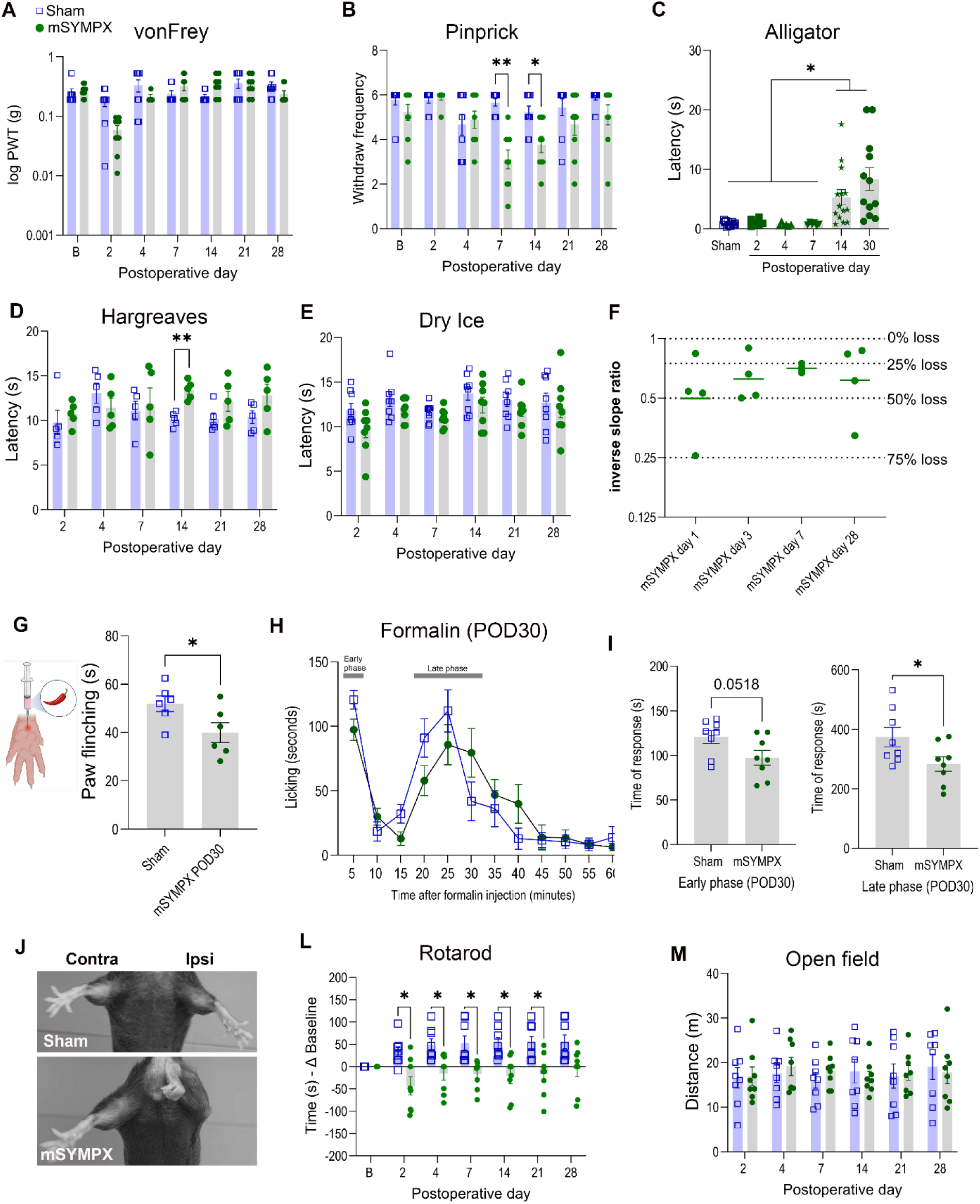
Assessment of sensory and motor functions in mice with targeted sympathetic denervation of the lumbar DRGs. **(A)** Time course of the von Frey test comparing mechanical sensitivity in sham and mSYMPX mice (n=8). **(B)** Pinprick responses differed significantly at POD 7 and 14 (n=9). **(C)** Alligator clip test showed increased withdrawal latency in mSYMPX animals at POD 14 and 30 (n=6-14). **(D)** Hargreaves test revealed altered thermal sensitivity at POD 14 (n=5). **(E)** Dry ice test showed no difference in cold sensitivity between groups (n=8). **(F)** The number of L4 DRG neurons responding to von Frey poking of the hindpaw were reduced after mSYMPX (n=4). The inverse ratio was plotted (ipsi slope/contra slope). Ratio <1 corresponds to fewer cells responding on ipsilateral side. Slope computed separately for each animal and each line indicates mean (for log of ratio values). **(G)** Capsaicin paw injection elicited fewer flinches in mSYMPX compared to sham at POD 30 (n=6). **(H)** Time course of formalin-evoked pain behavior (n=8) and **(I)** cumulative duration of formalin-evoked responses demonstrated significant group differences (n=8). **(J)** Gait analysis showed altered paw positioning in mSYMPX mice. **(L)** Rotarod performance indicated persistent motor impairment in the mSYMPX group (n=8). **(M)** Open field testing revealed no differences in locomotor activity between groups (n=8). Data are presented as mean ± SEM. Statistical analysis for **A, B, D, E, L, M:** two-way ANOVA with Sidak’s post hoc test. *p < 0.05, **p < 0.01 vs. sham; **C:** one-way ANOVA with Tukey’s multiple comparison test, *p < 0.05; **G, I:** unpaired student t-test.

Although changes in mechanical sensitivity were detected in the pinprick test, both von Frey and the pinprick tests activate very few receptive fields due to the small contact areas. To address the limitations, we utilized the alligator test to apply pressure force to the entire hind paw. This behavioral assay is like the tail clip assay described by Ranade et al.(*40*) and measures the response time to an alligator clip applied to a mouse’s hind paw. To ensure consistent placement, the clips were marked at the center and were positioned across the middle of the hind paws in all mice tested. A response was recorded when the mice exhibited awareness of the clip through behaviors such as paw withdrawal, flinching or shaking. The data was plotted as response latency for each mouse (Fig. 3C). The alligator test can activate multiple nerve fibers, offering a more comprehensive indicator of neuron loss compared to the von Frey or pinprick tests, which engage only a small number of neurons. Increased response latency was observed on POD14, indicating a significant alteration in sensory perception or pain response in mice. This increased latency persisted through POD30, suggesting that the altered sensory perception due to the loss of neurons may be long-lasting.

Increased latency of withdrawal response to heat stimulation was observed in the paws ipsilateral to the mSYMPX on POD14 as assessed using the Hargreaves test (Fig. 3D). This finding indicates a significant alteration in the thermal pain response in the affected limb due to the loss of small nociceptive neurons. However, no difference in cold sensitivity was detected in animals with mSYMPX on any time point tested between POD2 and POD 28 (Fig. 3E).

### Decreased number of neurons evoked by mechanical stimulation of the ipsilateral hind paw in mSYMPX mice

With the findings from the behavioral assessment, we proceeded to use the *in vivo* whole DRG calcium imaging, a technique successfully employed in our prior studies (*17, 18*), to quantify the number of neurons activated by mechanical stimulation of the hindpaw in anesthetized animals with and without sympathetic denervation of the DRGs.

To visualize neuronal activity *in vivo*, we expressed the ultrasensitive calcium indicator GCaMP6s in DRG neurons by crossing Pirt-Cre mice with Rosa26-loxP-STOP-loxP-GCaMP6s mice. After exposing the lumbar DRGs, neuronal activity was visualized using a fluorescence microscope equipped with a high-speed sensitive camera, capable of capturing rapid changes in calcium signals. We applied nylon filaments with ascending forces ranging from 1.4 to 15 grams to the hindpaw. In normal mice with intact sympathetic innervation of the DRGs, we observed a pronounced linear increase in the number of neurons firing as filament size increased. In contrast, mice with sympathetic denervation of the lumbar DRGs exhibited a significant decrease in neuronal firing in response to higher filament strengths (Fig. 3F and Suppl. Fig. 2). These findings are consistent with the loss of sensory neurons resulting from sympathetic denervation-induced cell death.

### Sympathetic denervation altered responses in acute pain models

We next tested whether mSYMPX-induced neuronal cell death would alter responses in acute pain models. To address this question, we utilized both the capsaicin and formalin models, which are extensively used in the field of pain research for their ability to induce and measure acute pain responses.

In the capsaicin model, we injected 20 µl of capsaicin (200 pmol) into the heel of the hind paw ipsilateral to the mSYMPX. Following the injection, we recorded and counted the number of paw flinches over a period of 5 minutes. The results revealed a significant decrease in the number of paw flinches in mice that had undergone sympathetic denervation of the lumbar DRGs compared to controls (Fig. 3G). The observed reduction in paw flinches in response to capsaicin injection can be attributed to the loss of CGRP-positive peptidergic sensory neurons, which are known to express the capsaicin-receptor TRPV1 and play a crucial role in nociceptive signaling.

Formalin injections into the hind paw evoked the expected biphasic nociceptive response, characterized by flinching and licking behaviors. As anticipated, mice displayed a pronounced early neurogenic phase followed by a later inflammatory phase (Fig. 3 H-I). Quantification of cumulative flinches revealed that mSYMPX-treated animals exhibited markedly reduced nociceptive behaviors compared with sham controls. This reduction was evident in both the early phase (0–5 minutes) and in the cumulative response to late phase (0–25 minutes), indicating that targeted sympathetic denervation attenuates formalin-induced pain.

### Impaired motor functions in mice with sympathetic denervation of the DRGs

While mice with sympathetic denervation appeared generally normal and showed no obvious differences compared to sham-control animals, noticeable abnormalities were evident when the mice were handled by holding the base of their tails. Specifically, when a mouse without mSYMPX was lifted, its hind limbs extended with toes spread apart, demonstrating a typical instinctive reaction. In contrast, mSYMPX mice displayed an altered posture: the ipsilateral hind limbs were drawn close to the abdomen, and their toes were curled inward towards the sole of the paw. This change in posture was first observed on POD4 and persisted through POD30, indicating a long-lasting alteration following sympathectomy (Fig. 3J).

We further evaluated motor skills using the rotarod test and the open field test. The rotarod test assesses motor coordination and balance by measuring the duration an animal can remain on a rotating rod. Starting from POD2, mSYMPX mice demonstrated increased slips and falls from the rod compared to sham-operated animals, suggesting a decline in strength and coordination. Changes in motor coordination persisted throughout the testing period between POD2 and 21 (Fig. 3L). In the open field test, no significant differences in travel distance were observed between mSYMPX and sham-control mice at any time point from POD2 to POD28 (Fig. 3M). These findings indicate that sympathetic denervation of the DRGs leads to notable impairments in motor function, particularly in coordination and posture.

### Caspase-3 activation and apoptotic protein expression are changed following sympathetic denervation of the DRG

The caspase-3 assay measures the activity of this key executioner enzyme in apoptosis indicating the onset of this programmed cell death, often before visible changes in cell morphology are evident (*41*). We employed this assay to investigate the underlying mechanisms of sympathectomy-induced cell death in the DRG. Our results revealed a significant increase in apoptotic cells in the DRGs following mSYMPX, as evidenced by a higher number of caspase-3-positive neurons on POD1 (Fig. 4A-B).

**Figure 4.**
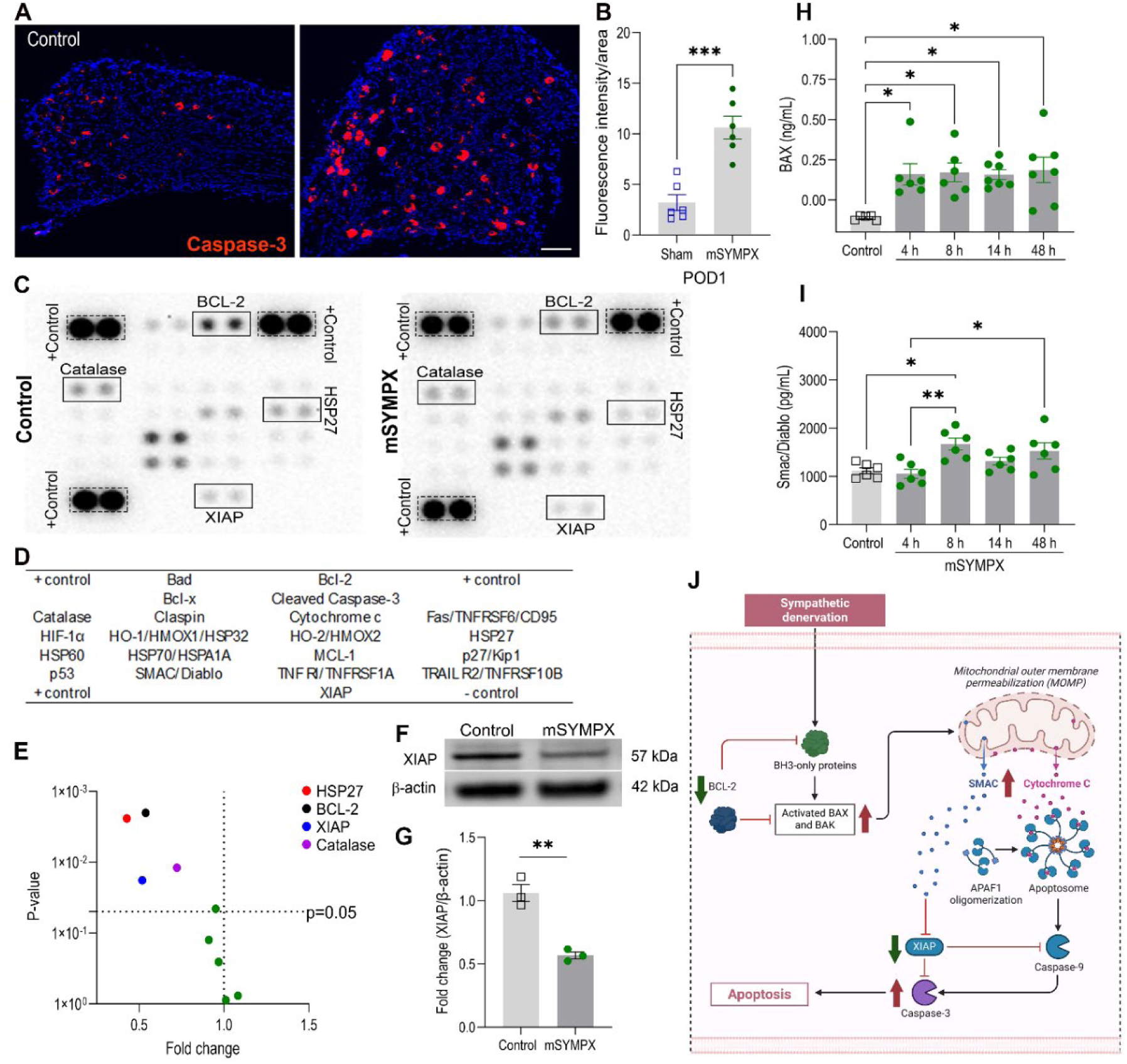
mSYMPX triggers apoptotic pathways in DRG neurons. **(A)** Representative immunofluorescence images showing elevated cleaved caspase-3 expression in L4 DRG 16 hours after mSYMPX. **(B)** Quantification of cleaved caspase-3 fluorescence intensity normalized to the cellular area confirmed increased expression in the mSYMPX group (unpaired t-test: ***p < 0.001 vs. sham, n=6). **(C)** Proteome profiling identified differentially expressed proteins between sham and mSYMPX groups. **(D)** Illustration of the cytokine array containing 21 different antibodies with duplicates. The array also contains three positive control (PC) proteins with strong signals in three corners of the membrane (for each membrane was used n=3). **(E)** Volcano plot highlighting downregulation of anti-apoptotic proteins in DRG tissues with mSYMPX. **(F-G)** Western blot analysis showing XIAP downregulation at POD2, with band quantification confirming reduced expression level (unpaired t-test: *p < 0.05 vs. sham, n=3). (H) ELISA analysis demonstrated upregulation of the pro-apoptotic proteins BAX (n=5-7) and Smac/Diablo **(I)** (one-way ANOVA with Tukey’s post hoc test; *p < 0.05 vs. control, n=6). **(J)** apoptosis pathway summary.

We further assessed the relative levels of mouse apoptosis-related proteins using a membrane-based antibody array (Fig. 4C-D), which allows simultaneous detection and quantification of multiple proteins in a sample. Among the 21 apoptosis-related proteins measured from three mice per group, four anti-apoptotic proteins (Heat Shock Protein 27 (HSP-27), B-cell lymphoma 2 (BCL-2), X-linked Inhibitor of Apoptosis Protein (XIAP), catalase) were downregulated on POD2 (Fig. 4C-E). The XIAP protein downregulation was confirmed by Western Blot (Fig. 4F-G). Using ELISA assay, we also revealed that the BAX apoptotic protein in the DRGs began to rise 4 hours after mSYMPX and continued to be elevated after 48 hours (Fig. 4H). Smac/Diablo peaked at 8 hours and continued to be upregulated after 48 hours (Fig. 4I). Smac antagonizes inhibitor-of-apoptosis proteins, as demonstrated by the observed decrease in XIAP in the antibody array experiment and Western Blot (Fig. 4F-G). This is the likely cause of the higher number of caspase-3 positive neurons as shown in the apoptosis pathway summary (Fig. 4J).

### Macrophage activation in the DRGs with sympathetic denervation

We previously reported that the sympathetic nervous system plays a crucial role in maintaining immune homeostasis (*16*). To explore whether macrophages in the DRG are involved in mSYMPX-induced cell death, we examined mRNA expression of macrophage signals. We observed altered expression on POD 1 in the DRG with mSYMPX compared to naïve mice with no nerve injury or inflammation. Among the genes we tested, *Cd68, Cx3cr1*and *Ccr2* were increased, while no change occurred for Iba1. By POD7, all macrophage related genes examined were increased in the mSYMPX DRGs (Fig. 5A-C).

**Figure 5.**
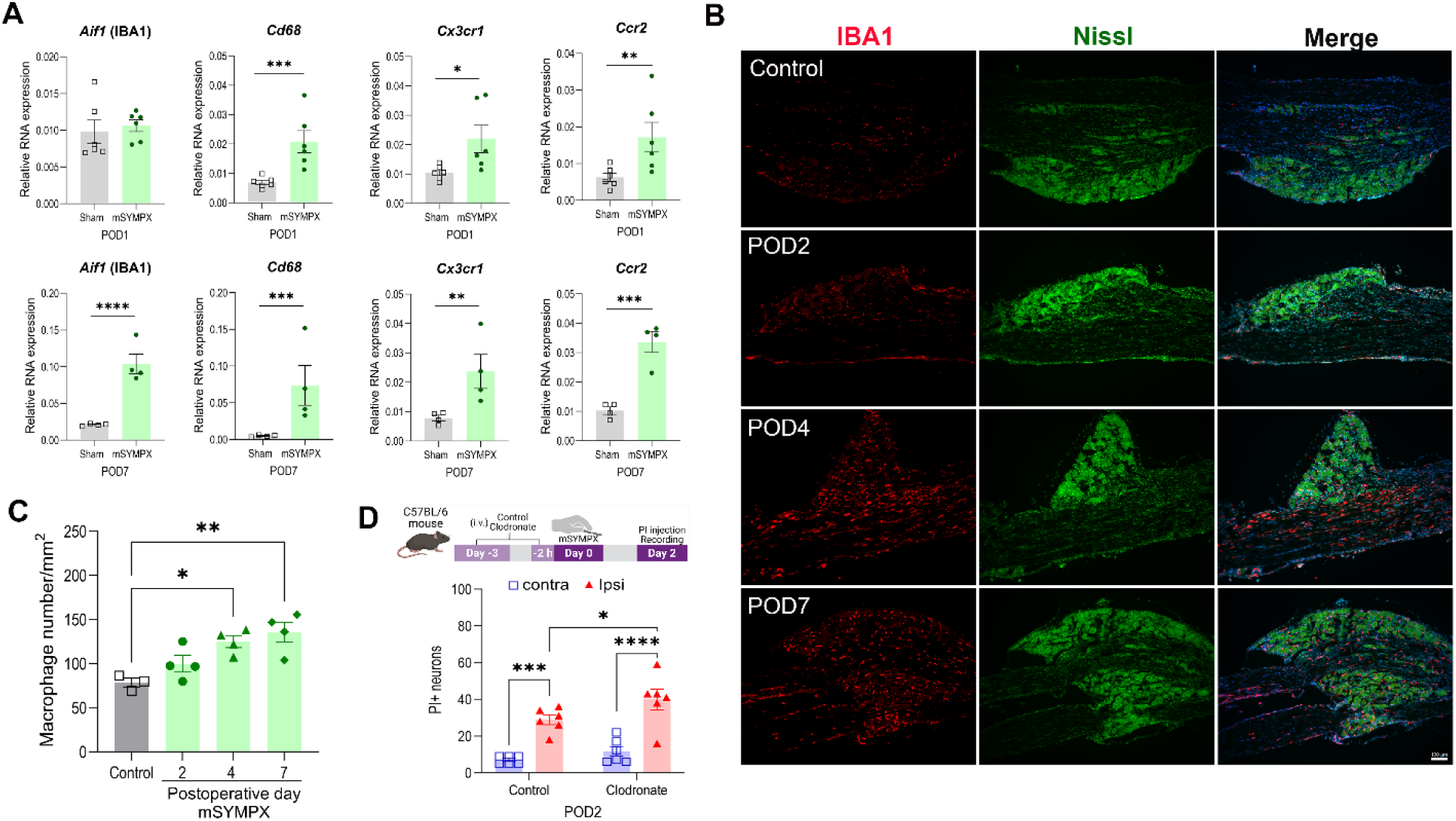
Macrophage activation in the DRG with sympathetic denervation. **(A)** Gene expression profile of different macrophage markers (*Aif1, Cd68, Cx3cr1, Ccr2*) showing upregulation after mSYMPX at POD1 (top panel) and POD7 (bottom panel). Data was analyzed by unpaired t-test, *p < 0.05, **p < 0.01, ***p < 0.001, ****p < 0.0001 (n=6). **(B)** Representative immunofluorescence images showing increased macrophage presence in DRG at POD2, 4, and 7. Macrophages were stained with IBA1 (red) and counterstained with Nissl (green) and DAPI (blue). Scale bar = 100 µm **(C)** Quantification of macrophage density (cells/area) confirmed progressive increase at POD 2, 4, and 7 (N=4). **(D)** Macrophage depletion prior to sympathetic denervation failed to prevent neuronal loss and even increased cell loss measured on POD2 after mSYMPX (two-way ANOVA with Sidak’s post hoc test: *p < 0.05, ***p < 0.001, ***p < 0.0001 vs. sham, n=2). Data are presented as mean ± SEM.

We then conducted an experiment comparing the degree of cell death in mice subjected to sympathetic denervation, with and without macrophage depletion to understand the role of macrophage activation in mSYMPX-induced cell death (Fig. 5D). Macrophage depletion was achieved by intravenous (i.v.) administration of liposomal clodronate 3 days prior to surgery, followed by a second dose on the day of surgery before the mSYMPX procedure (Suppl. Fig. 3) (*42*). We assessed cell death by mSYMPX using PI staining of the DRGs contralateral and ipsilateral to the mSYMPX. Control group mice received unilateral mSYMPX but no macrophage depletion using i.v. administration of a liposome vehicle solution.

The results demonstrated that i.v. administration of liposomal clodronate did not cause cell death in the DRGs contralateral to mSYMPX. In the sympathetic denervated DRG, however, macrophage depletion caused additional cell death measured by the number of PI+ neurons compared to the mSYMPX DRG without macrophage depletion (Fig. 5D), suggesting that macrophage activation following surgical sympathectomy may be an immune response triggered to protect cells from further damage.

### Sympathetic denervation-induced cell death is prevented by prolonged NE supplementation

We next examined noradrenergic signaling in mSYMPX-induced cell death, since norepinephrine is the main neurotransmitter released from sympathetic nerve endings.

We first assessed the expression of adrenergic receptors in dorsal root ganglia (DRG) from control and sympathectomized animals. Multiple adrenergic receptor subtypes, including both α-and β-adrenoceptors, are expressed in neuronal and non-neuronal populations within the DRG. In naïve animals, *Adra2c* and *Adrb1* are predominantly expressed in DRG neurons, whereas *Adrb2* is mainly expressed in macrophages and satellite glial cells (*43*). Following targeted sympathetic denervation, *Adra2c* expression was rapidly downregulated at postoperative day 1 (POD1) and remained suppressed through POD7 (Fig. 6A–B). Similarly, *Adrb1* expression decreased by POD7, coinciding with the downregulation of neuronal markers at the same point (Fig. 2C and Suppl. Fig. 1). In contrast, *Adrb2* expression showed a trend towards-upregulation at POD1 and further increased by POD7, which is consistent with the expansion of macrophage/immune cell populations observed at this stage (Fig. 5A–C). These transcriptional changes therefore reflect the dynamic regulation of adrenergic receptors across surviving neuronal and non-neuronal cells in the DRG following sympathectomy.

**Figure 6.**
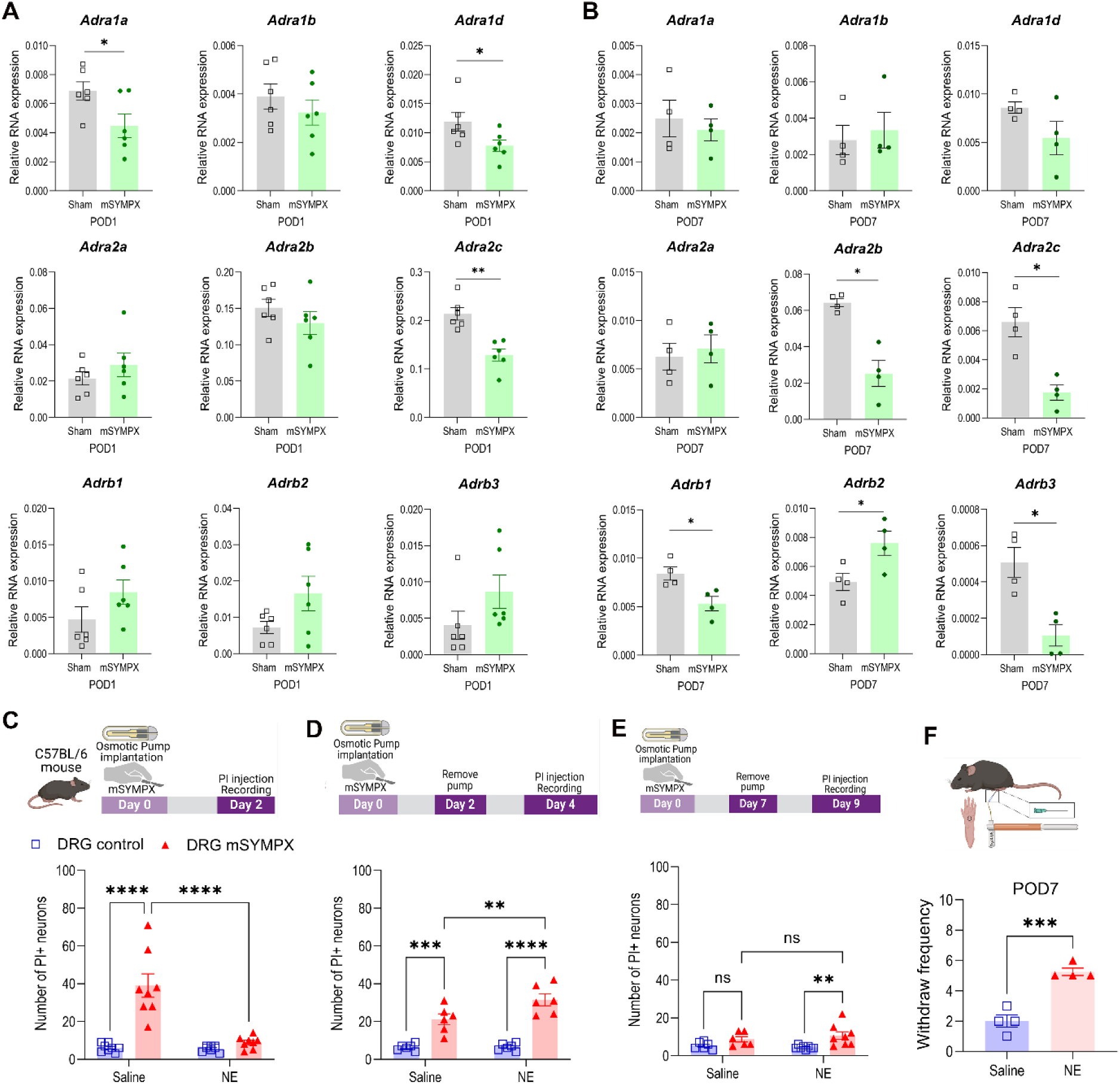
Sympathetic denervation–induced cell death is prevented or rescued by norepinephrine (NE) supplementation. **(A)** Adrenergic receptor expressions were downregulated at POD1 following mSYMPX (n=6). **(B)** Further downregulation of adrenergic receptors was observed at POD7 (n=4). **(C)** NE supplementation at the time of surgery prevented mSYMPX-induced cell death, as shown by in vivo imaging of PI⁺ neurons in the DRG at POD2 (n=8). **(D)** NE supplementation initiated at surgery but discontinued at POD2 failed to prevent cell death when imaging was performed at POD4 (two days after pump removal) (n=6). **(E)** NE supplementation maintained for 7 days and removed at POD7, followed by imaging at POD9, continued to protect against cell death (n=8). **(F)** Continuous NE supplementation for 7 days also restored behavioral responses to pinprick stimulation (n=4). **A, B, F:** Data analyzed by unpaired t-test, *p < 0.05, **p < 0.01, ***p < 0.001; **C-E:** Two-way ANOVA with Sidak’s post hoc test, **p < 0.01, ***p < 0.001, ****p < 0.0001.

Finally, we aimed to determine whether sympathetic denervation-induced cell death could be prevented or rescued by supplementing NE. In the first experiment, NE or vehicle was delivered via an osmotic pump (3.5 µg/Kg/min) implanted subcutaneously 4 hours before surgery to reach a stable NE supplementation when mSYMPX surgery was performed to avoid any gap of supplementation. The osmotic pump provided a steady release of NE for up to 3 days. *In vivo* imaging of the DRG was performed on POD2 to assess the number of PI+ neurons (Fig. 6C).

In the second experiment, mice received NE or vehicle supplement as above, but the pump was removed on POD2, with imaging performed on POD4, two days after pump removal (Fig. 6D). In the third experiment, mice received NE or vehicle for a full 7 day post-surgery, with the pump removed on POD7 to eliminate potential NE supply at that time; imaging was performed two days later (POD9) (Fig. 6E).

In the first experiment, where continuous NE supply was maintained and imaging recording occurred on POD2, cell death by mSYMPX was effectively prevented, as indicated by PI+ neuron count comparable to those in DRGs without mSYMPX. However, in the experiment with the NE pump removed on POD2, cell death increased to slightly above the level seen in the control group without NE supplementation. Conversely, when the NE pump was removed at a later time point on POD7, the level of cell death detected in the mSYMPX DRG remained low, similar to that observed in the mSYMPX DRG lacking NE supplementation. These results suggest that continuous NE supply is crucial for more than two days following sympathetic denervation to effectively prevent cell death.

To verify the effectiveness of norepinephrine (NE) supplementation up to POD7, prior to pump removal, animals were evaluated using the pinprick test. NE-supplemented animals exhibited a significantly higher frequency of withdrawal responses compared with the control group (mSYMPX + saline pump) (Fig. 6F). These responses were like those observed in naïve animals (Fig. 3B).

## Discussion

This study reveals a previously unrecognized physiological role of the sympathetic signaling in supporting the survival and functional integrity of sensory neurons within the DRG. Our findings show that targeted sympathetic denervation by localized microsympathectomy of the lumbar DRGs leads to considerable neuronal apoptosis, resulting in reduced spinal projections, decreased sensory innervation in peripheral tissues and changes in sensory/motor behaviors.

These results indicate that sympathetic nerves deliver essential trophic support through NE, which is vital for maintaining the viability and proper functioning of sensory neurons.

A key novel insight of this work is the identification of sympathetic dependence as a fundamental aspect of sensory neuron health, extending beyond its conventional roles in autonomic regulation, stress responses, and immune regulation. The significant neuron loss following sympathetic denervation, coupled with the protective effect of NE supplementation, indicates that NE acts as a neuroprotective agent in the DRG. This aligns with existing literature highlighting NE’s roles in neuroprotection across various neural systems, including its mitigation of oxidative stress, attenuation of neuroinflammation, and promotion of survival pathways such as PI3K/Akt signaling (*44–48*). The continuous NE supply appears critical within an early post-injury window (∼2 days), emphasizing the importance of timing for potential therapeutic intervention.

Furthermore, our results reveal that sympathetic denervation elicits an immune response involving immediate macrophage activation within the DRG. The increase in macrophage-related gene expression and the exacerbation of neuronal death upon macrophage depletion suggest an innate immune component that can be both protective and detrimental, depending on context (*49*). The activation of macrophages following sympathetic denervation-induced sensory neuron death may serve a protective role by helping to prevent further neuronal loss, in addition to clearing debris and initiating repair processes. This is supported by the sequence of events observed after local sympathectomy: immediate cell death occurs shortly after surgical intervention, which is then followed by macrophage activation and recruitment. The neuronal death appears to subside within 3–4 days post-surgery, suggesting that macrophage activation may contribute to halting the ongoing apoptotic process initiated by the loss of sympathetic innervation.

The behavioral assays show that the loss of sensory neurons manifests as reduced responses to mechanical and thermal stimuli, evidenced by increased response latencies in pinprick, alligator, Hargreaves, and in vivo calcium imaging assays. Importantly, the observation that withdrawal responses to intense stimuli decrease after sympathetic denervation supports the notion that sympathetic signaling contributes to intact sensory functions to detect nociceptive inputs Conversely, clinical interventions like sympathetic blockades are sometimes employed to alleviate pain syndromes such as CRPS by local immune regulation (*16*). Our data suggests that at least in normal or early post-injury states, sympathetic input is essential for sensory neuron survival, indicating that sympathetic blockade’s efficacy may depend on the context, timing, and underlying pathology, possibly by disrupting maladaptive sympathetic-sensory interactions in chronic pain states.

The failed detection in mechanical sensitivity with von Frey assay is surprising despite the reduced number of neurons responding to paw stimulation with similar nylon filaments. Although the precise cause of this inconsistency remains unknown, one possible explanation is that the imaging experiments were conducted in anesthetized animals, thereby lacking the normal integration and processing typically provided by the central nervous system. This study, therefore, underscored a significant limitation associated with relying on a single behavioral assay for the precise evaluation of pain sensitivity. A more informative and comprehensive assessment would necessitate a multidisciplinary strategy, combining multiple testing methodologies.

The alterations in gene expression profiles further corroborate the neurodegenerative effects of sympathetic removal. Downregulation of neuronal markers and neuropeptides such as CGRP and TRK-A (p75) post-sympathectomy indicate that sympathetic denervation of the DRG not only causes cell death, but surviving neurons also undergo phenotypic changes, potentially impairing their sensory functions. These changes could result in persistent sensory deficits and altered pain perception observed behaviorally.

Our findings further highlight the critical role of NE in preventing apoptosis, as continuous NE delivery through an osmotic pump markedly decreases neuronal death. The timing underscores the importance of early supportive interventions to maintain neuronal integrity following injury or sympathetic denervation. Considering NE’s broad neuroprotective functions such as mitigating oxidative stress and neuroinflammation (*44, 45, 47, 48*), therapeutic approaches that sustain or emulate sympathetic signaling may prove beneficial in conditions characterized by sensory neuron degeneration including neurotrauma and neurodegenerative disorders.

While this study emphasizes the role of sympathetic input in neuronal survival, further research is necessary to identify specific cellular targets of NE within the DRG, including the potential roles of satellite glia. Based on our findings, we believe that the loss of sympathetic NE removes tonic adrenergic trophic support to DRG neurons, which is mediated by β/α receptor–linked cAMP/PKA and PI3K/Akt survival pathways. This disruption leads to mitochondrial dysfunction characterized by shifts in BCL-2 family balance, disturbed Ca^2+^ homeostasis, and increased ROS/ER stress. Such mitochondrial disturbances promote outer membrane permeabilization and result in the release or increased expression of Smac/DIABLO. Our data support this, showing that Smac levels in the DRG increase immediately after surgical sympathectomy. Smac antagonizes inhibitor-of-apoptosis proteins, as reflected by the observed decrease in XIAP in our protein array and western blot. This facilitates the activation of caspase-3, leading to the execution of the intrinsic apoptotic pathway.

In conclusion, our findings demonstrate that targeted sympathetic denervation induces profound structural and functional remodeling within the lumbar DRG. The loss of sympathetic fibers surrounding sensory neurons eliminates adrenergic input and trophic support, thereby altering neuronal survival pathways, gene expression, and excitability, while potentially reducing aberrant sympathetic–sensory coupling implicated in pain states. Concurrently, the absence of sympathetic vascular regulation increases local blood flow and vascular permeability, promoting immune cell infiltration. Resident macrophages and satellite glial cells may respond to these changes by adopting activated phenotypes and releasing both pro- and anti-inflammatory mediators, which in turn modulate neuronal function. Together, these findings indicate that sympathetic denervation reshapes the DRG microenvironment at neuronal and immune levels, with outcomes that may either mitigate or exacerbate pain signaling depending on the physiological and pathological context. These insights open new avenues for therapeutic approaches, ranging from sympathetic modulation to targeted neurotrophic support to prevent or reverse sensory neuron loss and manage pain conditions. Recognizing the sympathetic nervous system not merely as an effector of autonomic control but as an essential guardian of sensory neuron integrity underscores its potential as a target in neuroprotection and chronic pain management.

## Materials and Methods

### Experimental Design

We investigated the impact of targeted sympathetic denervation of lumbar DRG on sensory neuron survival, function, and local immune responses. Our objectives were to determine: 1) if sympathetic denervation triggers neuronal death and degeneration; 2) the contribution of macrophages to these outcomes; and 3) whether continuous norepinephrine (NE) supplementation mitigates sympathetic denervation-induced changes.

To address these questions, we utilized the surgical mSYMPX procedure in mice, selectively removing the gray rami entering the L3–L4 DRG. Experimental groups included sham surgery, mSYMPX alone, mSYMPX with macrophage depletion using clodronate liposomes, and mSYMPX with continuous NE supplementation delivered via osmotic minipumps. Outcome measures, assessed at defined time points and in both sexes, comprised: behavioral assays for sensory function; in vivo calcium imaging (GCaMP6s/Pirt-cre mice) for neuronal activity; and molecular and histological analyses for neuronal death (PI staining, apoptosis arrays, ELISA, Western Blot), macrophage recruitment (immunohistochemistry), gene expression (qPCR), and structural changes within the DRG, spinal projections, and skin.

### Animals

The study was approved by the Institutional Animal Care and Use Committee of the University of Cincinnati. Male and female C57BL/6J (8–12 weeks, stock number 000664) and Ai14 floxed/conditional reporter mice (Ai14, stock number 007914) were used in this study and obtained from Jackson Laboratory, Bar Harbor, ME. Rosa26-lox-stop-lox GCaMP6s and Pirt-cre mice were obtained from the laboratory of Dr. Xinzhong Dong. These two strains were crossed to express the calcium indicator GCaMP6s in sensory neurons. Ai14 and Pirt-cre mice were crossed to obtain a reporter mice with sensory neurons labeled. All experiments included animals of both sexes in approximately equal numbers. We did not observe sex differences in any of the outcomes measured, so sex was not included as a variable. All efforts were made to minimize animal suffering, reduce the number of animals used, and use alternatives to in vivo techniques, in accordance with the International Association for the Study of Pain guidelines and the National Institutes of Health Office of Laboratory Animal Welfare Guide for the Care and Use of Laboratory Animals, and adhered to animal welfare guidelines established by the University of Cincinnati Institutional Animal Care and Use Committee who approved the experimental protocols used.

### Reagents

Capsaicin (M2028, Sigma-Aldrich), Formalin (SF100, Fisher Chemical), clodronate liposomes and control liposomes (CP-005-005, Liposoma, The Netherlands), noradrenaline bitartrate (5169, Tocris), 0.9% saline solution (50-238-8187, Teknova), micro-osmotic pump (model 1003D and 1007D, Alzet), propidium iodide (P1304MP, Invitrogen), proteome profiler mouse apoptosis array (ARY031, R&D Systems), mouse BAX ELISA kit (NBP2-69937, Novus), and mouse SMAC ELISA kit (AB246534, Abcam).

### Localized microsympathecthomy (mSYMPX) for targeted sympathetic denervation (TSD) of the DRG

Microsympathectomy was performed as originally described in rats (*16*) and adapted for mice (*15*). Briefly, the proximal L3 and L4 spinal nerves and transverse processes on one side were exposed. The spinal nerves (ventral rami) were visualized and freed from surrounding tissue. The gray rami entering the L3 and L4 spinal nerves were identified ventrally, near their junction with the DRG at the intervertebral foramen. At this site, where each gray ramus merges into the spinal nerve, the gray rami and surrounding connective tissue were gently dissected away from adjacent blood vessels and then severed from the spinal nerve. Approximately 0.5 mm of the gray ramus was further removed to create a gap. This procedure was performed on both L3 and L4 gray rami. Sham controls received similar exposure of the spinal nerves, but their gray rami were not cut.

### PI staining

Propidium Iodide (PI; label for dying cells) was injected intravenously via superficial femoral veins. The mouse was sacrificed 30 min later and the lumbar DRGs (L3/L4) were removed and placed in the recording chamber as an *ex vivo* live whole mount preparation. PI stains the nuclei only of cells with a disrupted plasma membrane. Neurons with the nucleus labeled were considered positive for cell death.

### Macrophage/Monocyte depletion

Liposome-encapsulated clodronate was used to deplete phagocytic macrophages. Clodronate liposomes (15ml/kg) were injected (i.p.) 3 days and 2 hours before the mSYMPX surgery (*15, 50*).

### Continuous supplementation of norepinephrine (NE)

At the indicated time points, an osmotic minipump (model 1003D, Alzet) was subcutaneously implanted through a small incision, under sterile conditions and isoflurane anesthesia, in accordance with the manufacturer’s instructions. The minipump was prefilled with either norepinephrine (NE+) or saline as a vehicle control (NE-). Norepinephrine concentration was adjusted to deliver a dosage of 3.5 µg/kg/min, based on studies indicating that higher dosages of 10 and 35 µg/kg/min resulted in increased variability or adverse effects such as hypertension and mortality (*51*).

### Behavioral tests

#### von Frey test

Mechanical sensitivity of the hindpaw was tested using von Frey filaments with the up-and-down method (*52*). Experimenters were blinded to surgical procedures and treatments. Briefly, mice were first acclimatized to the room for 1 hour, followed by another 1-hour acclimatization period in individual clear Plexiglas boxes on an elevated wire mesh platform to facilitate access to the plantar surface of their hindpaws. Subsequently, a series of von Frey filaments (0.02, 0.04, 0.07, 0.16, 0.4, 0.6, 1.0, 1.4, 2.0 and 4 g; Stoelting Co., Wood Dale, IL) were applied perpendicular to the plantar surface of the hindpaw. Testing commenced with the application of the 0.6 g filament. A positive response was defined as a clear paw withdrawal or shaking. If a positive response occurred, the next lower filament was applied; conversely, if a negative response occurred, the next higher filament was applied. The threshold was determined following 6 stimuli and analyzed as the logarithmic transforms of the von Frey force, as described by Mills (*53*).

#### Pinprick test

Mice were acclimated in von Frey chambers for 1 hour. A 1-gram (bending force) nylon filament with a fine needle (27-G, BD Biosciences) attached at the end, was applied to the ipsilateral hind paw. The needle had a non-sharp tip to prevent skin penetration (*54*). Six trials per mouse were performed with at least 2 minutes intervals between each trial. Paw withdrawal was scored as a response and reported as total number of trials.

#### Paw pressure test (Alligator clip)

The paw pressure test was employed to assess response thresholds to mechanical pressure stimulation. This method was adapted from published reports (*55, 56*). Mice were gently restrained, and a flat alligator clip, delivering constant pressure, was applied to the mid-plantar surface of the ipsilateral hindpaw. The pressure was maintained until a paw withdrawal attempt was observed, and the time to respond, or latency, was recorded for each mouse. A cutoff of 30 seconds was imposed to prevent tissue damage.

#### Hargreaves test

Heat sensitivity in mice was assessed using the Plantar Test apparatus (IITC Life Science), which measures paw withdrawal latency in response to radiant heat stimulation. This assessment followed the Hargreaves method (*57*). A radiant heat source was directed at the mid-plantar surface of the mouse hindpaw until withdrawal occurred. To prevent tissue damage, a cutoff time of 20 seconds was applied for each stimulation. Each mouse underwent 2-3 trials, with ten-minute intervals between each test. The average of these recordings was used for statistical analysis.

#### Dry ice assay

Cold sensitivity was assessed in mice after acclimation to the Plantar Test apparatus (IITC Life Science) – the same setup used for the Hargreaves assay. Mice were acclimated for 45 minutes before dry ice stimulation, following a previously described protocol (*58*). For stimulation, a tightly packed 10 mL syringe containing dry ice was placed under the hindpaw. To prevent tissue damage, a cutoff time of 20 seconds was applied for each stimulation. Paw withdrawal latency was measured using a stopwatch. Each mouse was tested 2-3 times, with 10-minute intervals between trials. The average of these recordings was used for analysis.

#### Open field test

Mouse motor skills were assessed using the open field test, a measure of spontaneous locomotion. For this test, each mouse was placed in a 40 cm × 40 cm × 40 cm Plexiglas arena and allowed to freely explore for 15 minutes. The mouse’s movements were recorded during this period, and the total distance covered was measured using AnyMaze (Stoelting).

#### Rotarod test

The Rotarod test, which measures forced locomotion, involved placing each mouse on a rotating rod (IITC Life Science). The latency to fall was then measured as the rotation speed accelerated from 5 to 35 rpm over a 3-minute period. Each mouse was tested three times, and the average of these recordings was used for statistical analysis.

#### Capsaicin responses

To assess the acute pain behavior induced by capsaicin (nocifensive behavior) after mSYMPX on Post-Operative Day 30 (POD30), mice were initially placed in recording chambers for 45 minutes. Following this habituation period, mice were removed from the chambers, and an intraplantar injection of capsaicin (200 pmol, diluted in PBS in a 20 µL volume) was administered into their paws (*59*). The mice were immediately returned to the recording chambers, and video recording began. All nocifensive behaviors (flinching and licking) were observed and recorded for 5 minutes.

#### Formalin test

Mice were habituated in a test chamber for at least 45 minutes prior to intraplantar formalin injection (20 µl, 2%) into the hindpaws. Following injection, mice were immediately returned to the chamber, and flinches were recorded for 1 hour. Formalin induces a characteristic biphasic nociceptive response in mice, manifested as behaviors such as flinching, paw lifting, and licking. These phases are defined as Phase 1 (early phase, reflecting neurogenic nociception) and Phase 2 (late phase, reflecting inflammatory nociception) (*60*). Cumulative flinches, specifically the sum of flinches from 0 to 5 minutes and from 0 to 25 minutes, were compared between sham and mSYMPX animals.

### In vivo calcium imaging in GCaMP6s/Pirt-cre mice

The lumbar DRGs were surgically exposed. Under pentobarbital anesthesia, mice were positioned on an Olympus BX fluorescent microscope stage and imaged using a fast EM-CCD camera with a 4x dry objective. Image acquisition occurred at a rate of 1.25-1.67 Hz. Following a baseline recording, von Frey filaments (1.4 - 15g) were applied to the heel region of the hindpaw. Images were then processed using SlideBook software to assemble time-lapses, which were subsequently analyzed to detect changes in green fluorescence indicative of GCaMP6s-mediated calcium signaling. The number of neurons exhibiting GCaMP6s transient signals was quantified.

### qPCR

The L3 and L4 lumbar DRGs were collected from sham and mSYMPX mice at selected time points. Mice were first anesthetized with pentobarbital sodium and transcardially perfused with PBS, followed by rapid removal of the DRGs. Total RNA was extracted using a Norgen RNA/Protein Purification Plus kit, and its amount and quality were assessed using a SimpliNano UV-Vis Spectrophotometer. This RNA was then converted into cDNA using a High-capacity RNA-to-cDNA kit (Thermo Fisher Scientific). Real-time quantitative PCR was performed on a QuantStudio™ 3 Real-Time PCR System using PowerUp SYBR™ Green Master Mix. All samples were analyzed at least in duplicate and normalized to the expression of hypoxanthine phosphoribosyltransferase (Hprt). All oligonucleotides are listed in supplementary Table 1.

### Western blot

Western blot analysis was performed on protein samples isolated after RNA extraction using a Norgen RNA/Protein Purification Plus kit, following manufacturer’s instructions. Protein concentrations were quantified using the Qubit protein assay (Cat. No. Q33211, Thermo Fisher Scientific). Samples (15 µg protein/lane) were separated by SDS-PAGE on a NuPage™ 4-12% gel (Cat. No. NP0321BOX) and subsequently transferred to a nitrocellulose membrane (Cat. No. LC2000, Invitrogen). After blocking with bovine serum albumin (BSA) for 1 hour, membranes were incubated overnight at 4 °C with X-linked inhibitor of apoptosis protein (XIAP) antibody (Rabbit, 1:500, Novus Biologicals, Cat. No. NBP220918). Subsequently, membranes were incubated with an anti-rabbit secondary antibody (1:2000, Cell Signaling, Cat. No. 7074S) for 1 hour. Detection was performed using SuperSignal™ West Dura Extended Duration Substrate (Thermo Fisher Scientific, Cat. No. 3475). Membranes were scanned using an iBright FL1000 (Invitrogen), and specific bands were identified based on their molecular sizes. Band intensities were quantified using CellSens Imaging software (Olympus) and normalized to β-actin (primary antibody: mouse, NB600-501, Novus Biologicals; secondary antibody for β-actin: anti-mouse, Cat. No. 7076S, Cell Signaling).

### Immunohistochemistry

For quantitative analysis, 12-μm thick sections were cut from mouse DRGs and 25-μm thick sections from spinal cords. Quantification was performed on 3-6 sections per animal, from a minimum of three animals per condition (i.e., at least 9 sections per timepoint). Images of both control and mSYMPX DRG and spinal cord sections were acquired from corresponding lumbar levels corresponding to projections from L3 and L4 DRG to the spinal cord. All images were taken with consistent settings using an inverted fluorescence phase-contrast microscope (BZX810, Keyence) at 10X, 20X, or 40X magnification and analyzed using CellSens Imaging software (Olympus). Negative controls, obtained by omitting the primary antibody, were used to establish background fluorescence thresholds and are not shown. All antibodies are listed in supplementary Table 2.

### Hematoxylin and Eosin (H&E) staining

Tissue samples were fixed in 10% neutral-buffered formalin and subsequently processed for paraffin embedding by the Integrated Pathology Research Facility at Cincinnati Children’s Hospital Medical Center. Five-micrometer sections were then cut, deparaffinized, rehydrated through graded ethanol, and stained with hematoxylin and eosin according to standard histological protocols. Finally, slides were dehydrated, cleared, and cover slipped for imaging.

### Mouse apoptosis array using proteome profiler

Two days post-surgery, animals were terminally anesthetized, and lumbar L3 and L4 DRGs were rapidly removed. These were then homogenized in a lysis buffer containing a cocktail of protease and phosphatase inhibitors. Protein concentration was subsequently measured using Qubit (Invitrogen). For protein array analysis, an array kit from R&D Systems (catalog number ARY031) was utilized, with all reactions performed according to the manufacturer’s protocol.

### Enzyme-linked immunosorbent assay (ELISA)

Quantification of SMAC and BAX protein levels was performed using commercially available sandwich ELISA kits (Mouse SMAC ELISA kit, Abcam, AB246534; Mouse BAX ELISA kit, Novus, NBP2-69937). All assays were conducted according to the manufacturers’ protocols. Absorbance was subsequently read at 450 nm, and protein concentrations were interpolated from standard curves generated with the recombinant SMAC and BAX proteins provided in the respective kits.

### Data analysis and statistics

Statistical analyses were conducted using Prism 10 (GraphPad Software, LLC, La Jolla, CA, USA). All quantitative data are presented as the mean ± standard error of the mean (SEM). For comparisons between two independent groups (e.g., sham vs. mSYMPX), statistical significance was determined using an unpaired Student’s t-test. When comparing two related groups or within-subject changes, a paired Student’s t-test was employed. For experiments involving more than two groups or factors, data was analyzed using one-way, two-way analysis of variance (ANOVA), or two-way repeated measure ANOVA, as appropriate. Following a significant ANOVA result, specific post-hoc tests (e.g., Tukey’s, Sidak’s) were applied for multiple comparisons, with the exact test used for each analysis detailed in the respective figure legends. The predefined criterion for statistical significance across all analyses was set at a p-value of < 0.05. Some illustrations in Fig. 3G, Fig. 4J, Fig. 5D, Fig. 6D-F and Fig. S2A for this study were created with assistance of BioRender (BioRender.com.).

## Acknowledgments

This publication was made possible, in part, using the Cincinnati Children’s Integrated Pathology Research Facility, RRID: SCR_022637.

## Author contributions

Conceptualization: JMZ, JAS, TB. Methodology: DDL, WX, ASP. Investigation: DDL, WX, ASP, JAS, TB, JMZ. Supervision: JMZ, JAS. Writing-original draft: JMZ, DDL. Writing-Review & editing: DDL, WX, ASP, JAS, TB, JMZ.

## Data and Materials availability

All strains and results are available upon request to JMZ. Supplementary Tables 1 and 2 provide, respectively, the complete oligonucleotide sequences and the antibodies used in these experiments.

