## Supplementary materials for "Sympathetic Controls Fate and Function of Adult Sensory Neurons"

Debora Denardin Lückemeyer *et al*.

**This file includes:**

Figs. S1 to S3
Tables S1 to S2

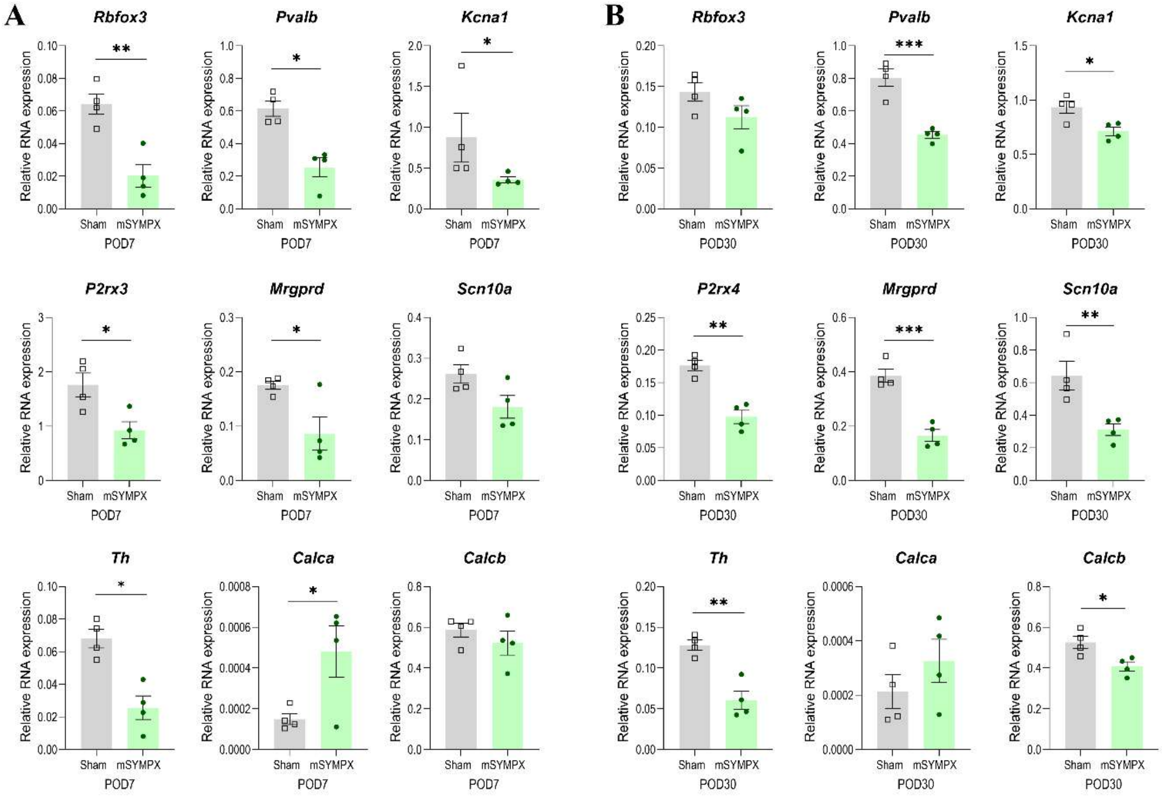


**Fig. S1. Changes in gene expression in the DRG with sympathetic denervation.**

Individual graphs for neuronal marker comparisons between sham and mSYMPX groups on POD7 **(A)** and POD30 **(B)** presented in Figure 2C. Data analyzed by unpaired t-test, $*p < 0.05$, $**p < 0.01$, $***p < 0.001$, n=4.

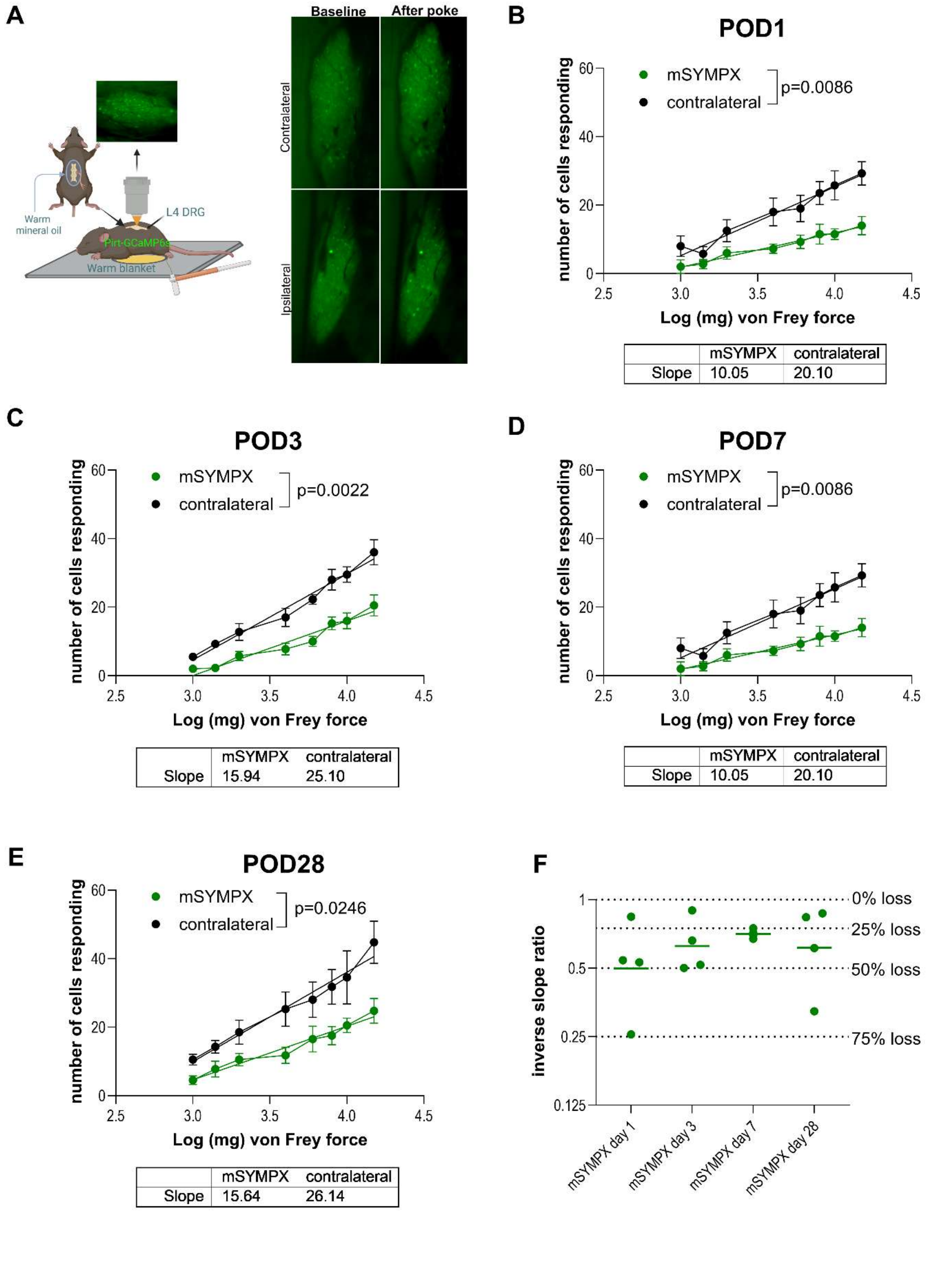
A
Baseline
After poke
Contralateral
Ipsilateral
Warm mineral oil
L4 DRG
Pirt-GCaMP6s
Warm blanket
B
POD1
mSYMPX
contralateral
p=0.0086
number of cells responding
Log (mg) von Frey force
Slope mSYMPX 10.05 contralateral 20.10
C
POD3
mSYMPX
contralateral
p=0.0022
number of cells responding
Log (mg) von Frey force
Slope mSYMPX 15.94 contralateral 25.10
D
POD7
mSYMPX
contralateral
p=0.0086
number of cells responding
Log (mg) von Frey force
Slope mSYMPX 10.05 contralateral 20.10
E
POD28
mSYMPX
contralateral
p=0.0246
number of cells responding
Log (mg) von Frey force
Slope mSYMPX 15.64 contralateral 26.14
F
inverse slope ratio
0% loss
25% loss
50% loss
75% loss
mSYMPX day 1
mSYMPX day 3
mSYMPX day 7
mSYMPX day 28

**Fig. S2. Altered neuronal responses in mice with mSYMPX.**

**(A)** Schematic of the *in vivo* recording setup used to quantify the number of L4 DRG neurons activated by mechanical stimulation of the paw with von Frey filaments. The number of neurons responding to paw poking was significantly reduced in the L4 DRG after mSYMPX compared with the contralateral DRG on POD1 **(B)**, POD3 **(C)**, POD7 **(D)**, and POD28 **(E)**. Reduced responses were observed for both innocuous and noxious mechanical stimuli. **(F)** Percentage loss of von Frey–responsive neurons, corresponding to data shown in panels **(B–E)**. Data analyzed by simple linear regression; *P* values for each comparison are reported within the corresponding panel.

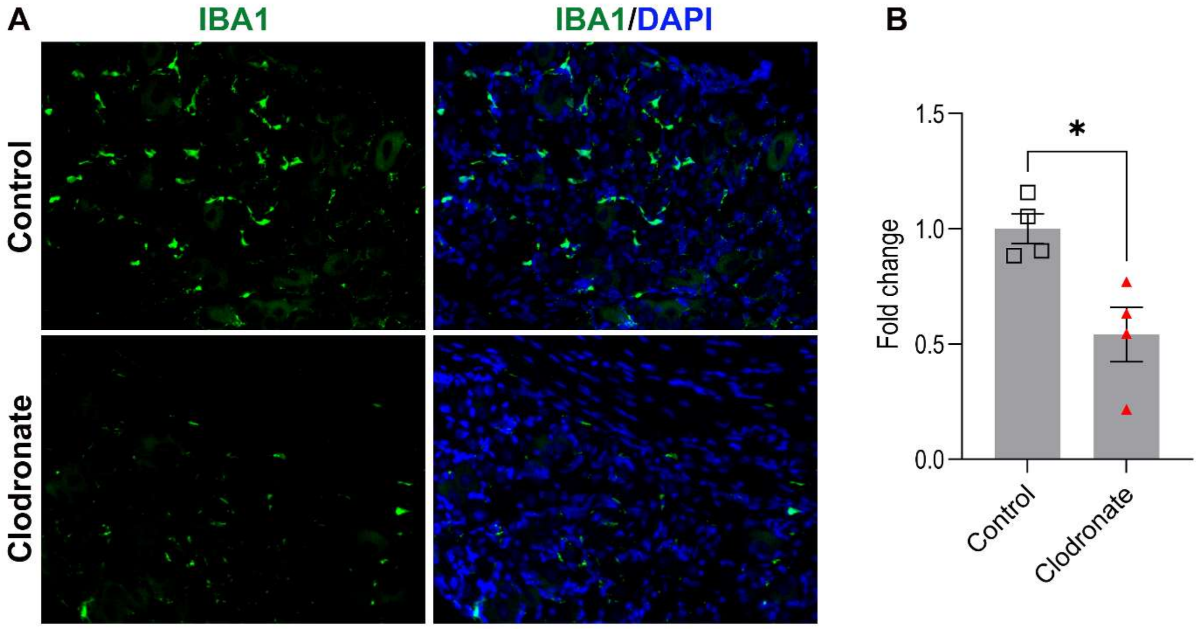


**Fig. S3: Macrophage depletion following clodronate liposome administration.**

**(A)** Representative immunofluorescence images of dorsal root ganglia (DRG) showing reduced macrophages after clodronate liposome treatment compared with control liposomes. Macrophages were labeled with IBA1 (green), and nuclei were counterstained with DAPI (blue). **(B)** Quantification of macrophage density expressed as fold change relative to control confirmed a significant reduction following clodronate liposome injection. Data are presented as mean ± SEM and were analyzed using an unpaired *t* test; $P < 0.05$.

**Table S1.**

Oligonucleotides for qPCR

| Protein | Gene | Forward (5’→3’) | Reverse (5’→3’) | Accession Number |
|---|---|---|---|---|
| NeuN | *Rbfox3* | CACCACTCTCTTGTCCGTTTGC | GGCTGAGCATATCTGTAAGCTGC | NM_001285437.1 |
| Parvalbumin | *Pvalb* | TGCAGACTCCTTCGACCACAAAA | GAGGAGAAGCCCTTCAGAATGG | NM_013645.4 |
| Kv1.1 | *Kcna1* | GAGTCGCACTTCTCCAGTATCC | CCCACGATCTTGCCTCCAATTG | NM_010595.3 |
| P2X3 | *P2rx3* | TCATCAACCGAGCCGTTCAGCT | ACTCTGTTGGCATAGCGTCCGA | NM_145526.3 |
| MrgprD | *Mrgprd* | GTCTTCCTCACCTGTTCTCTGC | CGAAGAGAAGCGTGATACGCAG | NM_203490.4 |
| Nav1.8 | *Scn10a* | ATGGAGGTCAGCCAGGACTACA | CTGTGAGGTTGTCCGCACTGAA | NM_009134.3 |
| TH | *Th* | TGCACACAGTACATCCGTCATGC | GCAAATGTGCGGTCAGCCAACA | NM_009377.2 |
| CGRP1 | *Calca* | GCACTGGTGCAGGACTATATGC | CTCAGATTCCCACACCGCTTAG | NM_007587.2 |
| CGRP2 | *Calcb* | GAGGAGCAAGAGACTAAGGGCT | GGCACAAAGTTGTCCTTCAGCAC | NM_054084.2 |
| IBA1 | *Aif1* | ATCAACAAGCAATTCCTCGATGA | CAGCATTCGCTTCAAGGACATA | NM_019467.4 |
| CD68 | *Cd68* | TGTCTGATCTTGCTAGGACCG | GAGAGTAACGGCCTTTTTGTGA | NM_001291058.1 |
| CX3CR1 | *Cx3cr1* | GAGTATGACGATTCTGCTGAGG | CAGACCGAACGTGAAGACGAG | NM_009987.4 |
| CCR2 | *Ccr2* | ATCCACGGCATACTATCAACATC | CAAGGCTCACCATCATCGTAG | NM_009915.2 |
| α-1A adrenergic receptor | *Adra1a* | GCCTCAAGACCGACAAGTCA | TCGGGAAGAAGGACCCAATG | NM_013461.5 |
| α-1B adrenergic receptor | *Adra1b* | GGCGGGAGTCATGAAGGAAA | GGAGAACAGGGAGCCAAGTG | NM_007416.4 |
| α-1D adrenergic receptor | *Adra1d* | GTGTCTTCGTCCTGTGCTGGTT | GCCAGAAGATGACCTTGAAGACG | NM_013460.5 |
| α-2A adrenergic receptor | *Adra2a* | CAGCCCCGTGTATAAAGCCA | GGACAGCAAGGGGAGGTAAC | NM_007417.5 |
| α-2B adrenergic receptor | *Adra2b* | GCAGAGAGCAGGGTGACAAT | AACCCACTTCCAGTTTGGGG | NM_009633.4 |
| α-2C adrenergic receptor | *Adra2c* | GCTTCAGGCAATGACCCTCT | AGCCCTAAGTTCACAGCCAC | NM_007418.3 |
| β-1 adrenergic receptor | *Adrb1* | CTCATCGTGGTGGGTAACGTG | ACACACAGCACATCTACCGAA | NM_007419.3 |
| β-2 adrenergic receptor | *Adrb2* | GGAACGACAGCGACTTCTT | GCCAGGACGATAACCGACAT | NM_007420.3 |
| β-3 adrenergic receptor | *Adrb3* | AGAAACGGCTCTCTGGCTTTG | TGGTTATGGTCTGTAGTCTCGG | NM_013462.3 |
| HPRT | *Hprt* | CTGGTGAAAAGGACCTCTCGAAG | CTGGTGAAAAGGACCTCTCGAAG | NM_013556.2 |

**Table S2:**
Antibodies for immunofluorescence and Western Blot

| Antibody/Dye | Catalog number | Vendor | Dilution | Host |
|---|---|---|---|---|
| Caspase-3 | AF835 | R&D Systems | 1:500 | Rabbit |
| CGRP | AB36001-1001 | Abcam | 1:500 | Goat |
| PGP9.5 | Ab108986 | Abcam | 1:500 | Rabbit |
| NF200 | N0142 | Sigma | 1:600 | Mouse |
| TH | P40101-150 | Pel-Freez | 1:500 | Rabbit |
| IBA1 | GTX100042 | Genetex | 1:500 | Rabbit |
| XIAP | NBP2-20918 | Novus Biologicals | 1:500 | Rabbit |
| β-actin | NB600-501 | Novus Biologicals | 1:2000 | Mouse |
| DAPI | D1306 | Invitrogen | 1:1000 | |
| IB4 Alexa Fluor™ 488 | I21411 | Invitrogen | 1:500 | |
| IB4 Alexa Fluor™ 647 | I32450 | Invitrogen | 1:500 | |
| NeuroTraceTM 500/525 green | N21480 | Invitrogen | 1:200 | |
| Neurotrace™ 435/455 blue | N21479 | Invitrogen | 1:200 | |
| Alexa Fluor™ 488 | 705-545-147 | Jackson Immuno Research | 1:300 | Goat |
| Alexa Fluor™ 488 | A21206 | Invitrogen | 1:1000 | Rabbit |
| Alexa Fluor™ 488 | A21202 | Invitrogen | 1:1000 | Mouse |
| Alexa 594 | A21207 | Invitrogen | 1:1000 | Rabbit |
| Alexa 594 | 715-585-151 | Jackson Immuno Research | 1:300 | Mouse |
| HRP-linked | 7074S | Cell Signaling | 1:2000 | Rabbit |
| HRP-linked | 7076S | Cell Signaling | 1:2000 | Mouse |
